# Expected patterns of local ancestry in a hybrid zone

**DOI:** 10.1101/389924

**Authors:** Joel Smith, Bret Payseur, John Novembre

## Abstract

The initial drivers of reproductive isolation between species are poorly characterized. In cases where partial reproductive isolation exists, genomic patterns of variation in hybrid zones may provide clues about the barriers to gene flow which arose first during the early stages of speciation. Purifying selection against incompatible substitutions that reduce hybrid fitness has the potential to distort local patterns of ancestry relative to background patterns across the genome. The magnitude and qualitative properties of this pattern are dependent on several factors including migration history and the relative fitnesses for different combinations of incompatible alleles. We present a model which may account for these factors and highlight the potential for its use in verifying the action of natural selection on candidate loci implicated in reducing hybrid fitness.

## 2 Introduction

A large fraction of research aiming to describe the process of speciation involves mapping genetic variants responsible for reproductive isolation. Despite its difficulty, this task has nevertheless been carried out for a number of cases in which the link between a reproductive isolating mechanism mapped in a laboratory setting and its effect on an individual’s fitness in nature is demonstrated [Schluter, 2009]. However, in many of these cases, reproductive isolation is already complete such that the initial cause of speciation cannot be attributed to any one locus or set of loci due to a lack of information regarding the order in which these isolating barriers arose [Turelli et al., 2014]. Hybrid zones present a convenient situation where reproductive isolation is incomplete. In these cases, the mechanisms of reproductive isolation are both fewer and more recently derived. Relative to scenarios with complete reproductive isolation, systems with ongoing hybridization may provide a more narrow set of candidate loci to consider as the initial drivers of speciation.

The next task would be to describe the mechanism by which the incompatible substitutions were fixed. Functional annotations for the implicated loci can yield some clues about the ecological context or genetic causes that resulted in these substitutions. A rigorously tested explanation would require that field experiments be carried out to establish their effect on fitness in nature [Schemske, 2000, Schemske and Bradshaw, 1999]. However, patterns of genomic variation can provide a complementary source of evidence for the action of natural selection on genetic variants which are relevant to a phenotype of interest [Tiffin and Ross-Ibarra, 2014]. The robustness of any given metric or model for the signature of natural selection depends on well-conceived theory that describes both the conditions under which the signature is detectable as well as any non-selective processes that can explain the pattern. This observational approach has been a driver of both theoretical and empirical research which aims to implicate loci responsible for genetic incompatibilities that decrease fitness among hybrids in nature [Barton, 1979, Barton and Hewitt, 1985, Endler, 1973, White, 1968].

Hybrid zones are thought to present a useful situation where the interaction between gene flow and natural selection can leave identifiable patterns associated with genetic incompatibilities in genomic data [Harrison and Larson, 2016, Payseur, 2010, Payseur and Rieseberg, 2016]. Historically, most work on this problem has relied on using differences in allele frequencies across the hybrid zone while ignoring patterns of linkage disequilibrium among neighboring sites [Barton and Hewitt, 1985]. More recently, increased access to sequencing technology has prompted the use of methods which can infer local ancestry across the genomes of admixed individuals [Gompert and Buerkle, 2013]. In this regard, population genetic inference has made a significant shift toward developing models which leverage this information for a variety of purposes. Several models aim to infer the migration history between genetically distinct populations using the length of ancestry tract lengths among admixed individuals [Gravel, 2012, Harris and Nielsen, 2013, Hellenthal et al., 2014, Liang and Nielsen, 2014, Loh et al., 2013, Patterson et al., 2012, Pool and Nielsen, 2009, Price et al., 2009, Sedghifar et al., 2015]. As the primary intention of these approaches has been to focus on populations within a species, there is a lack of work which aims to describe the effect of genetic incompatibilities which commonly arise between species after a prolonged period of geographic isolation.

Theory with formal treatment of genetic incompatibilities and ancestry tracts has been slow to accumulate, in large part due to the large parameter space of both migration histories and genetic architectures that may contribute to reduced fitness in hybrid individuals. As a result, forward simulations of whole chromosomes under differing migration and selection regimes have been used to describe some general patterns [Gompert et al., 2012, Hvala et al., 2018, Lindtke and Buerkle, 2015, Schumer and Brandvain, 2016]. In a few of these cases, the primary goal is to describe the conditions which may account for the heterogeneous patterns of genomic differentiation which have been widely observed across hybrid zones [Harrison and Larson, 2016]. For example, Gompert et al. [2012] focus on describing differences in both the number of contributing loci and the mechanism of their effect through either underdominance at single loci or two-locus epistasis. They also introduce a formalized approach to identify outlier loci responsible for reduced hybrid fitness using allele frequency clines across the genome. Lindtke and Buerkle [2015] pay particular attention to two-locus models of genetic incompatibilities and compare the relative efficiency with which different kinds of epistatic interactions can maintain genomic differentiation in a hybrid zone under both high and low migration.

In an effort to make use of ancestry tract lengths rather than allele frequencies at individual loci, Sedghifar et al. [2015] derive a null expectation for the length of ancestry tracts in a geographic context where distance from the contact zone of two genetically distinct populations is explicitly modeled. They then provide a likelihood function which they use to infer the age of the contact zone, or time at which admixture between the populations began. Sedghifar et al. [2016] extends this spatially-explicit framework further to model the mean ancestry tract length which is contiguous with an under-dominant locus.

Another approach that uses local ancestry inference to identify genetic incompatibilities relies on computing correlations in ancestry among pairs of loci in a hybrid zone [Schumer et al., 2014]. Schumer and Brandvain [2016] use simulation to demonstrate how selection against incompatible alleles at two loci can lead to a positive correlation in species ancestry at those loci. They find good power to identify these associations for genetic architectures that feature ubiquitous selection (see Figure 1d). The intuition for this pattern is that genotypes with the same ancestry at both loci are the only genotypes with high fitness, such that an over-representation of ancestry at those loci relative to background levels of linkage disequilibrium (LD) should lead to an identifiable signal. For genetic architectures that only feature strong selection against derived allele combinations, they find much less power to identify significant pairs.

**Fig 1.**
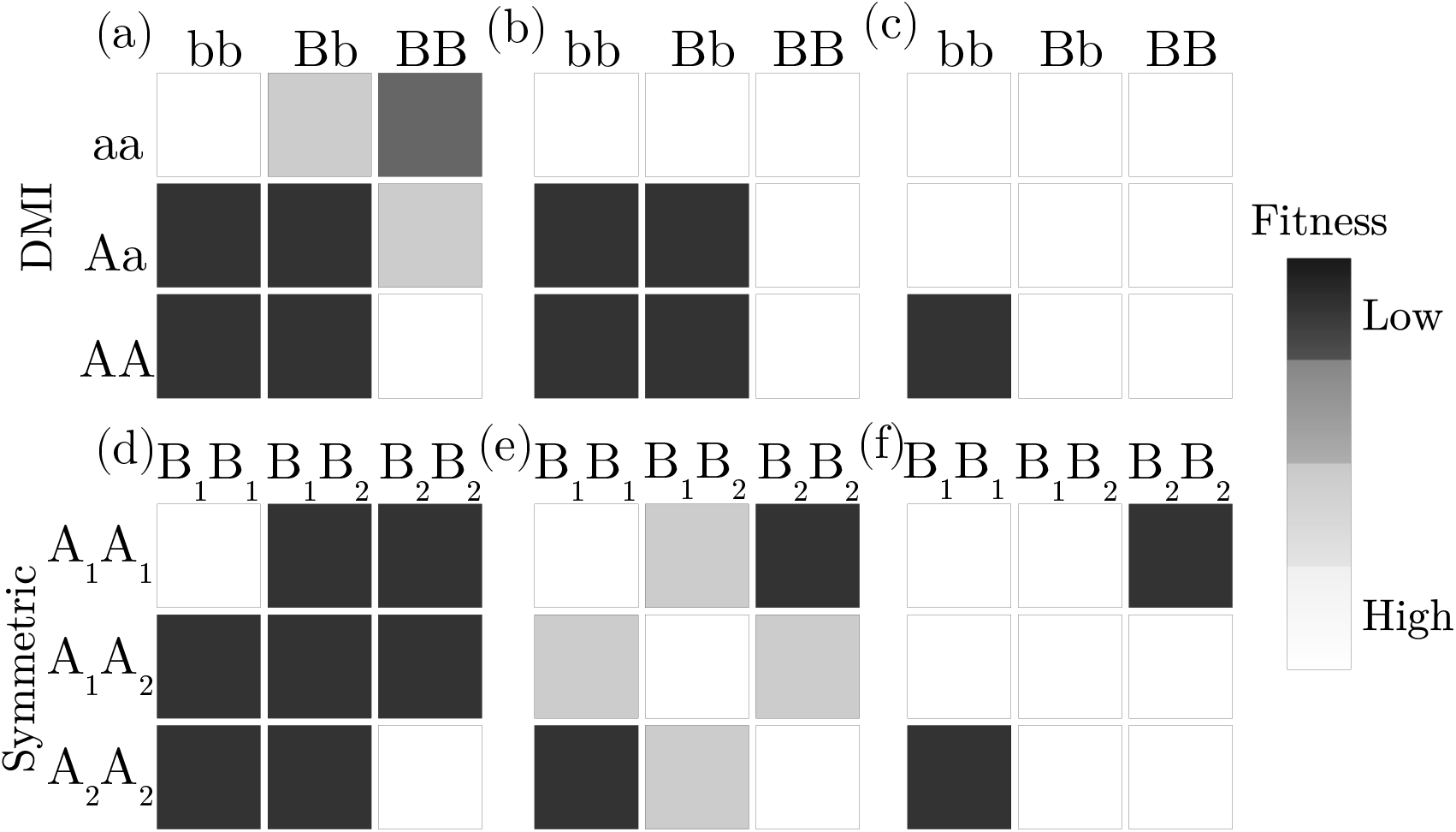
Two-locus fitness matrices for six models of genetic incompatibility. Each matrix includes the fitnesses of all possible two-locus genotypes where each locus is biallelic. Shaded boxes represent genotypes with a fitness cost that varies positively with the amount of shading. The top row of matrices are variations of the DMI model with the **aaBB** genotype representing the ancestral state and the bottom row shows variations of a symmetric incompatibility model. For both rows, the dominance effect of derived substitutions decreases from left to right.

The variety of approaches and data available to study this problem have prompted a few questions of where to proceed next. We first describe a few of the well-studied genetic architectures for two-locus genetic incompatibilities as well as others that have received less attention but which have also been identified in nature. We then present a model to compute the expected distribution of ancestry tract lengths around incompatibility loci.

### 2.1 Two-Locus Genetic Incompatibilities

The two-locus fitness matrix provides a useful representation of different genetic architectures which might contribute to genetic incompatibility between species (Figure 1 and Table 1). In addition to theoretical arguments and simulations, much of our current understanding for how relevant any of these genetic architectures might be in nature has been driven by genetic dissection of reproductive barrier phenotypes in the lab [White et al., 2011]. There are a number of empirical examples in a variety of species which have hinted at the potential importance of meiotic drive and neutral (or nearly neutral) causes for the fixation of incompatible substitutions [Maheshwari and Barbash, 2011, Presgraves, 2010, Sweigart and Willis, 2012]. While the precise combination of evolutionary forces which are responsible for incompatibility formation remain unknown, the evolution of incompatibilities in hybrid populations can be reasonably approximated with simple epistatis [Schumer et al., 2014].

**Table 1.** Genotype fitnesses for the DMI and symmetric incompatibility models. The first pairs of bold letters are DMI model genotypes and the genotypes in parentheses indicate the symmetric model. *s_a_* and *s_e_* denote the selection coefficient against the ancestral and incompatible alleles, respectively. *h_a_*, *h*_0_ and *h*_1_ denote the dominance effects of ancestral, double-heterozygotes and single-heterozygotes, respectively.

|  |  | **bb** (**B<sub>1</sub>B<sub>1</sub>**) | **Bb** (**B<sub>1</sub>B<sub>2</sub>**) | **BB** (**B<sub>2</sub>B<sub>2</sub>**) |
| --- | --- | --- | --- | --- |
| **aa** | (**A<sub>1</sub>A<sub>1</sub>**) | 1 | $1 - s_a h_a$ | $1 - s_a$ |
| **Aa** | (**A<sub>1</sub>A<sub>2</sub>**) | $1 - s_e h_1$ | $1 - s_e h_0$ | $1 - s_a h_a$ |
| **AA** | (**A<sub>2</sub>A<sub>2</sub>**) | $1 - s_e$ | $1 - s_e h_1$ | 1 |

The most well-known model is described in Dobzhansky [1937] in which alleles fix at two interacting loci among populations that are geographically isolated. The top row in Figure 1 shows a range of possible fitness matrices that might result from this scenario, also known as the Dobzhansky-Muller incompatibility model (DMI). If we denote the ancestral genotype as **aaBB** in all of these cases, then the derived genotypes before coming into secondary contact are **aabb** and **AABB**. We chose these example matrices to emphasize the diversity of fitness configurations that might result from this model. The fitness matrix in Figure 1a is an example where the the derived substitutions were fixed by positive selection, such that the ancestral genotype suffers a fitness cost. Figures 1a and 1b are examples where the derived alleles interact dominantly; whereas in Figure 1c, derived alleles interact recessively.

Lindtke and Buerkle [2015] draw attention to a different model of genetic incompatibility where allele substitutions occur at two loci in both populations leading to a symmetric pattern of fitnesses between the two derived genotypes **A_1_A_1_B_1_B_1_** and **A_2_A_2_B_2_B_2_** (Figure 1d, 1e, 1f). Their results suggest that this mechanism could provide a better explanation for the observed patterns of genetic differentiation that occur at extended genomic distances between species that hybridize [Harrison and Larson, 2016]. Regulatory interactions between a transcription factor encoded at one locus and the corresponding binding site at a second locus would be one scenario consistent with this model. Seehausen et al. [2014] note that this model could also be common in meiotic drive scenarios where a substitution that promotes biased transmission of a selfish genetic element at one locus is counteracted by a substitution at a second locus which restores unbiased inheritance. The bottom row in Figure 1 shows a range of possible fitness matrices under this model, where the left-most matrix results from dominant substitutions which interfere between haplotypes, and the right most matrix results from recessive substitutions. Simulated data in Figure 2 (using the software dfuse from Lindtke and Buerkle [2015]) illustrates the effect of the DMI model in Figure 1b where selection against derived alleles leads to a bias towards the ancestral genotype (**aaBB**) of recombined ancestries.

**Fig 2.**
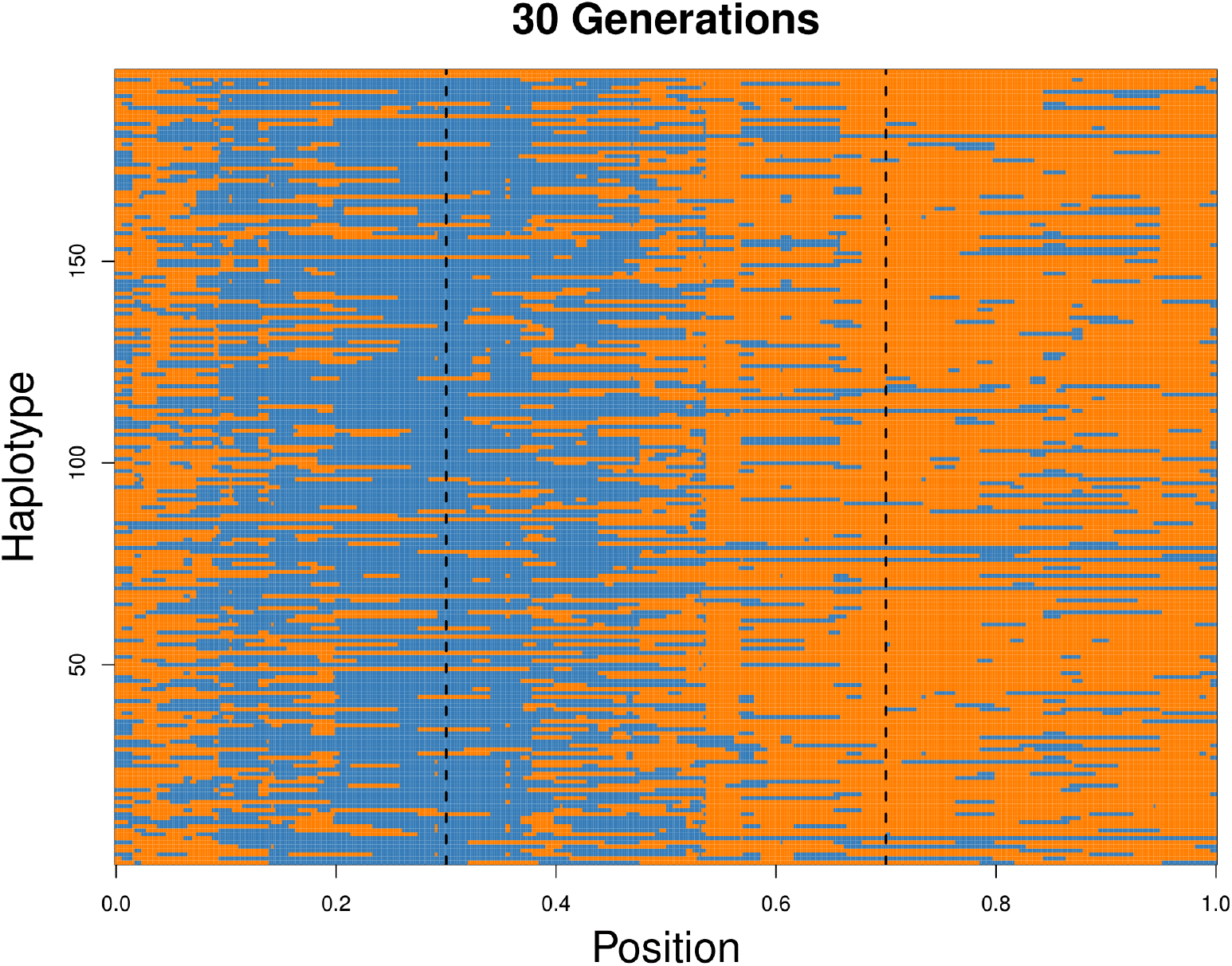
Haplotype data simulated using the software dfuse with the fitness matrix in Figure 1b. The forward-in-time simulation begins with two infinite source populations contributing equal fractions of ancestry (0.5) to a target population of 100 individuals 30 generations in the past. Each generation to the present follows a Wright-Fisher model, whereby both source populations contribute a fraction of individuals m to the target population. In this case *m* = 0.1. Recombination occurs uniformly along the chromosome at rate 1 crossover per chromosome per generation. After recombination, individuals are removed from the population according to a specified fitness matrix. The parameter values defined in Table 1 take the following values: *s_a_* = 0, *s_e_* = 0.9, *h*_1_ = 1, *h*_0_ = 1, and *h_a_* = 0. The incompatibility loci are indicated by the vertical dotted lines.

In the following section, we first review an approach taken by Gravel [2012] to model the distribution of ancestry tract lengths across the genomes of an admixed population. We then describe the framework for our own extension to this approach which aims to model the distribution of ancestry tract lengths that are contiguous with a locus undergoing epistatic interactions according to any of the incompatibility scenarios outlined above.

## 3 Model Description

### 3.1 Tract Length Distributions Under Neutral Admixture

Gravel [2012] defines a Markov chain along a chromosome with transition rates between both an ancestry state variable, *p*, and the time, *t*, at which ancestry *p* arrives in a hybrid population. Consider the demography of a sample up to the first hybridization event *T* generations ago, where each generation is labeled *s* ∊ {0,1,2,…,*T* − 1}. Let *m_p_*(*t*) denote the fraction of individuals in the target population replaced by individuals from source population *p* at time *t*. *m*(*t*) is the total fraction of individuals in the target population replaced by migrants in generation *t* where 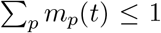. Moving along a chromosome from any point, the probability of encountering state (*p*,*t*) after a recombination event that occured at generation *τ* is

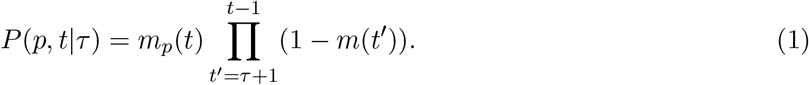

*τ* is uniformly distributed on (1, *t* − 1), so the discrete transition probabilites can be expressed as

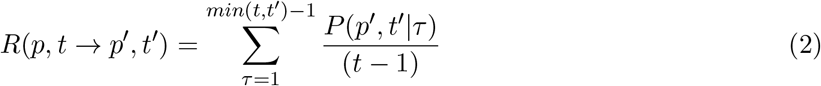

To get the continuous transition rate, one can multiply the discrete transition rate by the continuous overall transition rate *t* − 1. This follows from the fact that a recombination event occurs at each generation such that probability of observing an ancestry junction depends on the number of generations since admixture:

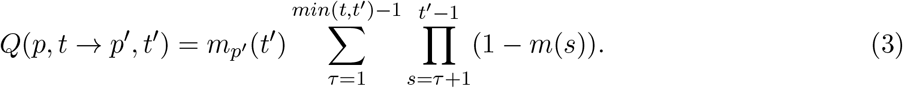

Using *Q*, one can compute the tract length distribution for a given ancestry. *Q* is first uniformized to adjust self-transition probabilities such that the total transition rate from each state is equal to the rate of the state with the highest transition rate, *Q*_0_ [Stewart, 1994]. One can then compute the distribution of the number of steps spent in a particular ancestry, {*b_n_*}_*n*=1,…,Λ_, up to a cutoff *Λ*, where 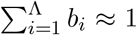 {*b_n_*}_*n*=1,…,Λ_ is computed by multiplying the state vector with the transition matrix for Λ iterations while recording the amount of probability absorbed by the non-*p* ancestries at each step. The Erlang distribution models the length of a trajectory, *l*, with *k* steps as:

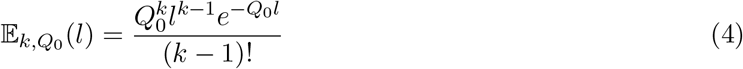

This leads to the tract length distribution:

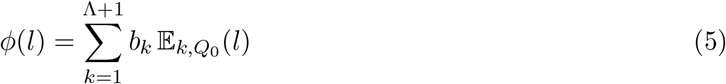

### 3.2 A Locus-Specific Tract Length Distribution With Selection

Equation 5 describes the length of tracts in a way that is not locus specific. We are interested in how the effects of purifying selection against alleles at two loci under negative selection, according to the incompatibility models described above, may skew the tract length distribution. More specifically, we want to model the distribution of ancestry tracts lengths that are contiguous with a negatively selected allele on a chromosome. In this case, the probability of observing a transition, or recombination event, depends on its recombination distance from the incompatibility loci of interest.

We define the number of basepairs between loci A and B to be *v* + *w* = *L*, where *v* is the number of basepairs from the A locus to the *v*th position and *w* is the number of basepairs from position *v* + 1 to *L* (Figure 3). We extend the transition matrix *Q* in equation (3) such that each value of *v* denotes a new *Q_v_* by multiplying each transition rate by the probability, 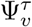, that an ancestry junction which arises at time *τ* at position *v* survives to the present:

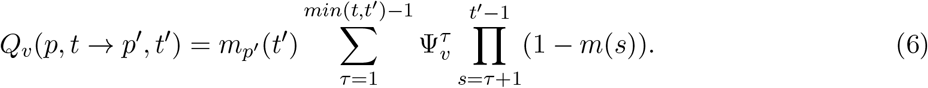

Equation 6 is computed as a function of the sequence of genotypic backgrounds the junction encounters each generation to the present. Using a two-allele model, let **A** and **a** refer to alternative alleles at the locus of interest, and alleles **B** and **b** refer to the second locus located at some distance away from the **A** locus. We can define a state space, *S*, of two-locus genotypes in which the junction can exist:

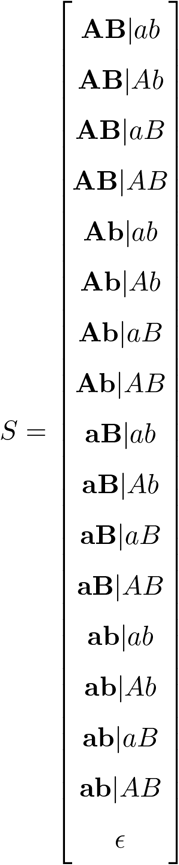

where the bold pair of alleles refers to the chromosome on which the junction resides. In cases where the interacting loci are on different chromosomes, the bold alleles refer to the genomic complement from which the junction is inherited.

**Fig 3.**
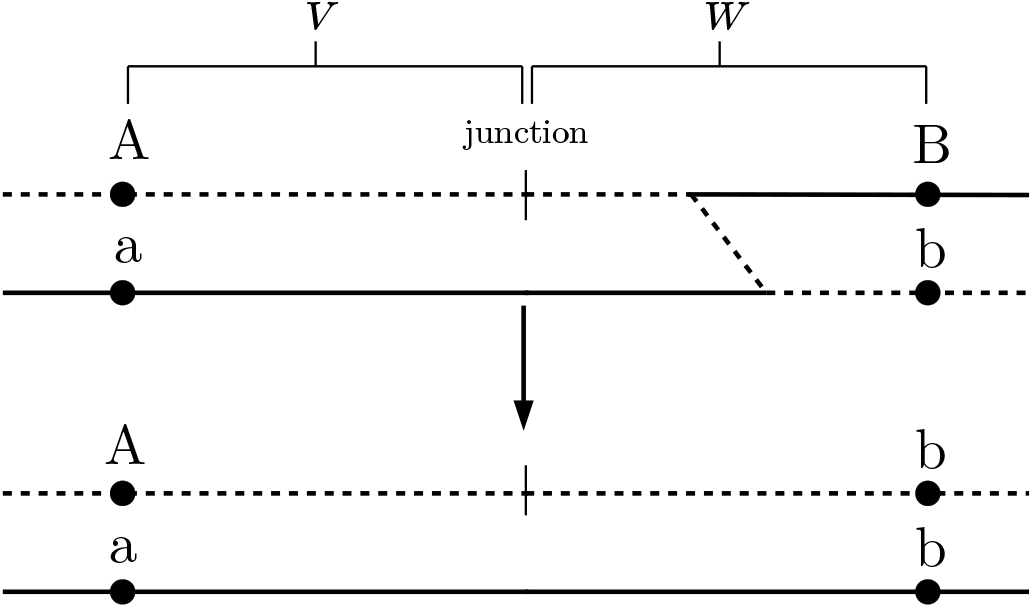
A visual description of the transition probability 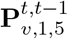. For the first state in *S*, **AB**|*ab*, the transition probability to state **Ab**|*ab*, is a product of the probability that the bold haplotype (**AB**) is chosen (0.5), a recombination event occurs between the junction and locus B, 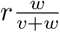, the recombined gamete gets paired with gamete *x*_4_ at time *t* − 1, and the individual with genotype **Ab**|*ab* survives, *ω*_14_.

Let 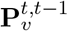 be a symmetric 17 × 17 transition matrix among the states in *S* from time *t* to *t* − 1 for a junction at the *v*th position, where 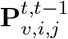 refers to the transition from state *i* to *j*. The first row in this matrix is shown below in Equation 7 (see Appendix A for rows 1-17). The transition probabilities in 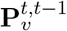 depend on the fitness of genotypes carrying the junction, *ω*, the recombination rate between the interacting loci, r, and the frequency of possible gametes with which to pair in the hybrid population at time 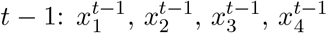. Let *x*_1_, *x*_2_, *x*_3_, *x*_4_ refer to the frequencies of gametes **AB**, **Ab**, **aB** and **ab**, respectively. Gamete frequencies are computed numerically by simulation [Gavrilets 1997, Appendix A.5]. Let *ω_i_* denote the marginal fitness of gamete *i* where *ω*_1_, *ω*_2_, *ω*_3_, *ω*_4_ refer to gametes **AB**, **Ab**, **aB** and **ab**, respectively. Let *ω_ij_* refer to the fitness of an individual with gametes *i* and *j*. Figure 3 provides some intuition for how the following transition probabilities in 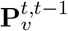 are computed.

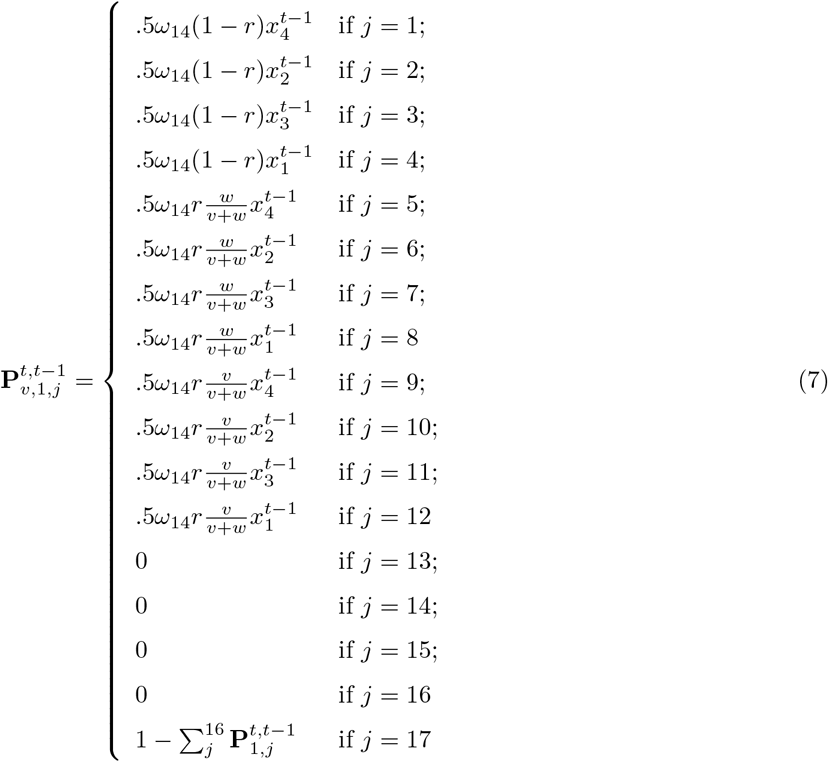

We can define the initial probabilities, 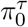, of a junction in each state when it occurs at a particular time *τ*. These probabilities will vary depending on the ancestry of interest for the tract length distribution. Conditional on a recombination event occurring between the two loci, the probability that the junction occurs at any particular position is uniform (1/*L*). If the ancestry of interest is that of the A allele, then

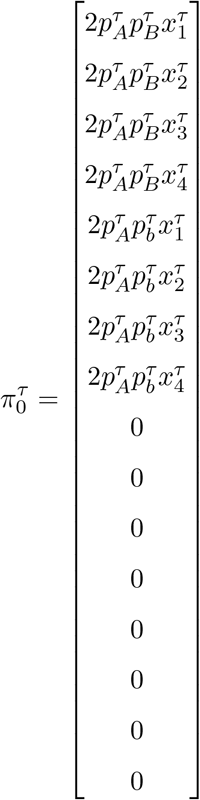

where 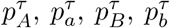 are the allele frequencies at time *τ*. The probability that the junction resides among each of the states after its origination at time *τ* to the present is

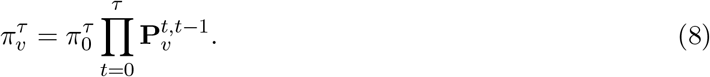

After defining the vector *η* = [0, 0, 0, 0, 0,0, 0, 0, 0, 0, 0, 0, 0, 0, 0, 0, 1], the survival probability of the junction is

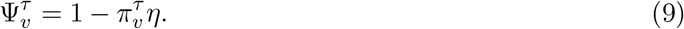

The transition matrix *Q_v_* can now be computed using Equation 6 for all values of *v* where *v* ∈ {1…*L*}. In contrast to the transition matrix *Q* defined in Equation 3, the set of transition matrices *Q_v_* are inhomogeneous over positions *v*. As a result, the uniformization technique outlined in Stewart [1994] does not apply. However, Andreychenko [2010] describes an approach to uniformize a time-inhomogeneous Markov chain which relies on partitioning the transition matrix into time-dependent and time-independent components. Whereas the time-homogeneous case relies on uniformizing by the constant transition rate of the state with the largest value, the time-inhomogeneous case relies on using the average rate of the state with the largest transition rate value. As before, the distribution for the number of steps in a trajectory, {*b_n_*}_*n*=1,…,Λ_, can be computed and used with Equations 4 and 5 to calculate the tract length distribution.

## 4 Discussion

The model presented above describes an approach which may prove useful in verifying the role of purifying selection against incompatible alleles in a hybrid zone. If shown to be robust under a reasonable set of demographic scenarios and genetic architectures for incompatibility, this model would provide an additional tool for testing the effects of selection on candidate loci which have been identified by QTL mapping of hybrid sterility or inviability traits [White et al., 2011]. This model could also be used to develop an independent test of loci identified by steep clines in allele frequency across a hybrid zone relative to the genomic background [Gompert et al., 2012]. We would like to emphasize the novelty and potential power of considering a genetic architecture of reproductive isolation for which there is strong empirical support (The Dobzhansky-Muller model) in the context of ancestry.

While any formal statements regarding the expected tract length distribution would require a full implementation of this model, we can make a few intuitive statements which follow from previous theory and simulation [Hvala et al., 2018, Lindtke and Buerkle, 2015]. In particular, Hvala et al. [2018] show that the number and density of ancestry junctions scales negatively with selection strength at incompatibility loci. This signature was further influenced by the genetic distance of junctions from the loci, the form of selection and dominance.

There are several challenges that remain before computing expected tract length distributions and performing inference on parameters of interest. In particular, computing *Q_v_* for a large set of positions may be difficult considering the repeated summation over products in Equation 6, the matrix multiplication required both for Equation 8, and computing {*b_n_*}_*n*=1,…,Λ_. While Gravel [2012] intended to model admixture events which occurred relatively recently, many hybrid zones of interest are likely to have formed more than 100 generations ago, which produces more computational burden given that the state space of *Q_v_* is 2*Tv*. However, it is likely that differences in the junction survival probability, 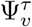, beyond some value of *τ* become negligible. The simplified two-locus, two-allele model that we consider is another effort to reduce the parameter space of genotype fitnesses that might result from higher-order epistasis of 3 or more loci.

Because 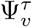 is dependent on hybrid zone gamete frequencies in a linear stepping-stone model, deviation from this simplifying assumption will most likely affect the results. The linear stepping-stone model which we borrow from Gavrilets [1997] can be generalized to any number of demes between the two infinite source populations. By implementing our model with this population structure, one could compute tract length distributions as a function of distance from the hybrid zone in a similar spirit to the more geography-explicit approach of Sedghifar et al. [2015, 2016]. The assumption of large population size is another assumption which could also be relaxed (see Appendix 1 in Gravel [2012]), as this will likely influence the rate at which ancestry junctions are fixed or lost from the population.

Aside from the challenges of model misspecification, performing inference will be particularly difficult considering the computational burden of computing the tract length distribution for a set of migration rates and fitness matrix parameters. Gravel [2012] uses a maximum-likelihood scheme to identify the set of parameters that best describe the magnitude and timing of migration events from a source into a target population. Given that our primary interest is to infer the effects of purifying selection, it may be more efficient to treat the migration history as a latent variable to be marginalized over using Markov chain Monte Carlo.

Despite these challenges, our framework for computing statistical properties of haplotypes in a hybrid zone represents one of only a few recent efforts which aim to exploit the combination of whole-genome sequencing and dense genotyping approaches that have emerged for non-model systems. In particular, this model is the only example that we know of for deriving locus-specific haplotype patterns under epistatis. Given the complexity of this problem, an alternative option may be to use simulation-based classification in a machine learning framework [Chan et al., 2018, Schrider and Kern, 2018, Sheehan and Song, 2016]. Rather than focusing on any one summary statistic, several summary statistics with potential relevance to purifying selection against genetic incompatibilities could be used simultaneously. Alternatively, Chan et al. [2018] describe another machine learning approach which could instead use genotype data directly.

Regardless of the methods used to identify genomic patterns of purifying selection against incompatibility loci, this effort represents one facet of the many lines of evidence necessary to identify and describe the causes of reproductive isolation between species.

## Acknowledgements

We thank Megan Frayer and John Hvala at The University of Wisconsin, Madison for their helpful discussions and insight. We also thank Yaniv Brandvain for additional advice and encouragement. Members of the Novembre, Stephens, and He labs provided useful feedback at an early stage. This work was funded by NSF grant DEB-1353737 to Bret Payseur, DEB-1353737 to Bret Payseur and John Novembre as well as NSF Graduate Research Fellowship and National Institute Of General Medical Sciences of the National Institutes of Health under award numbers DGE-1144082 and T32GM007197 to Joel Smith.

## Appendix A

The full transition matrix used to compute the junction survival probabilities in Equation 9:

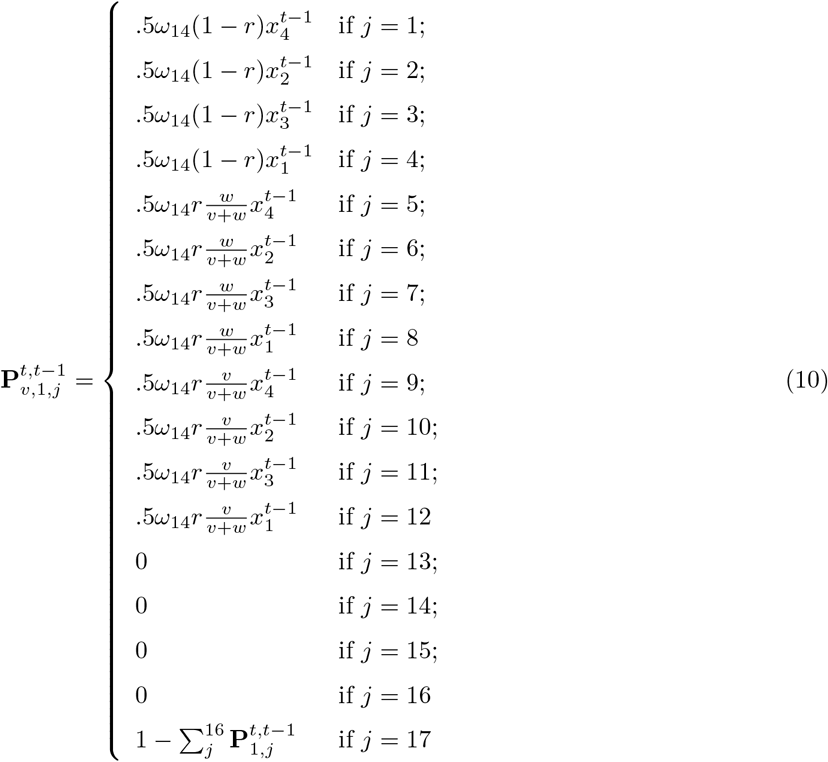

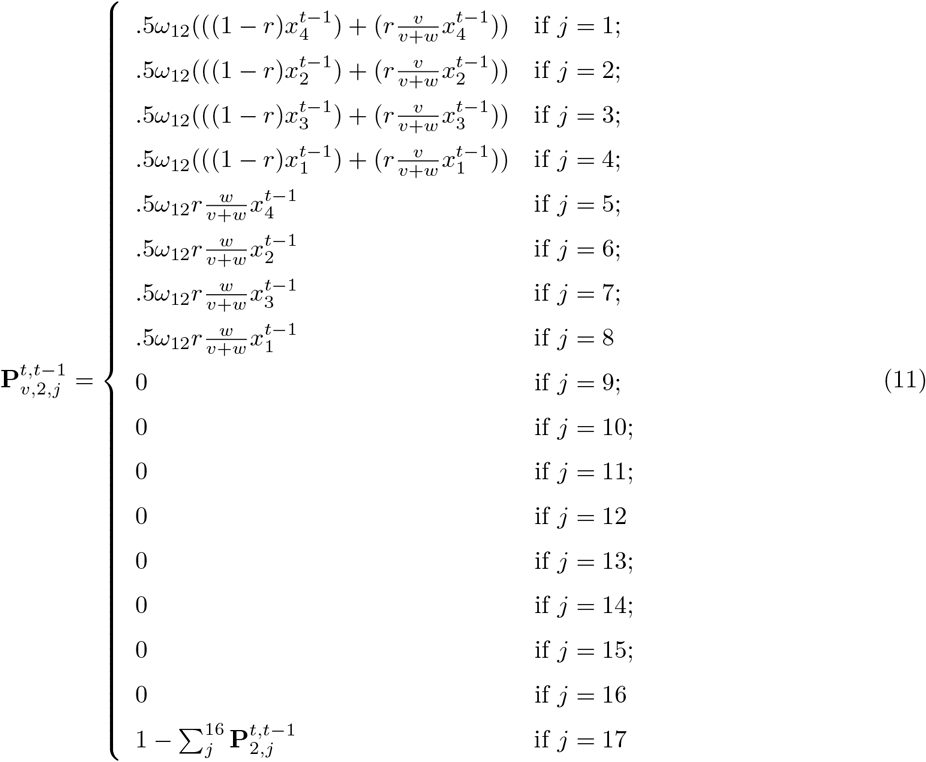

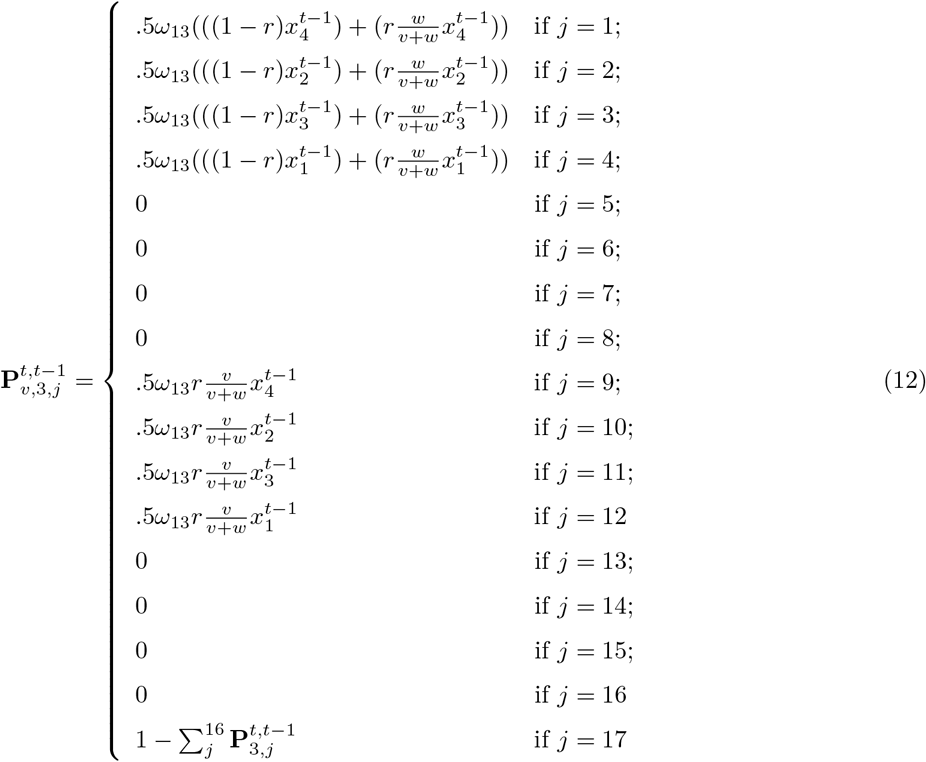

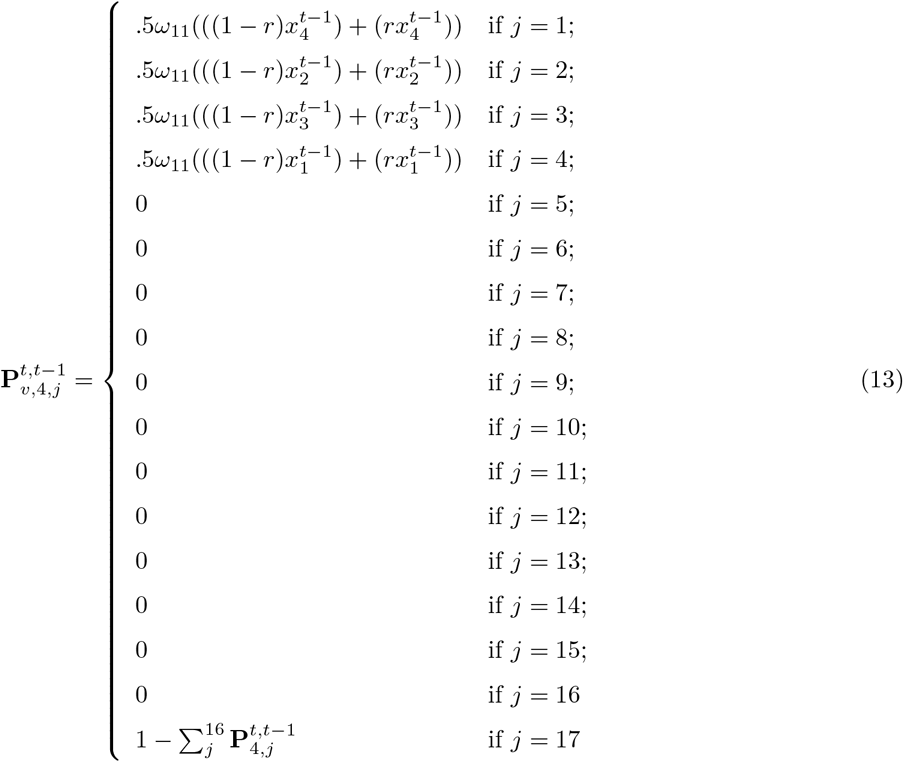

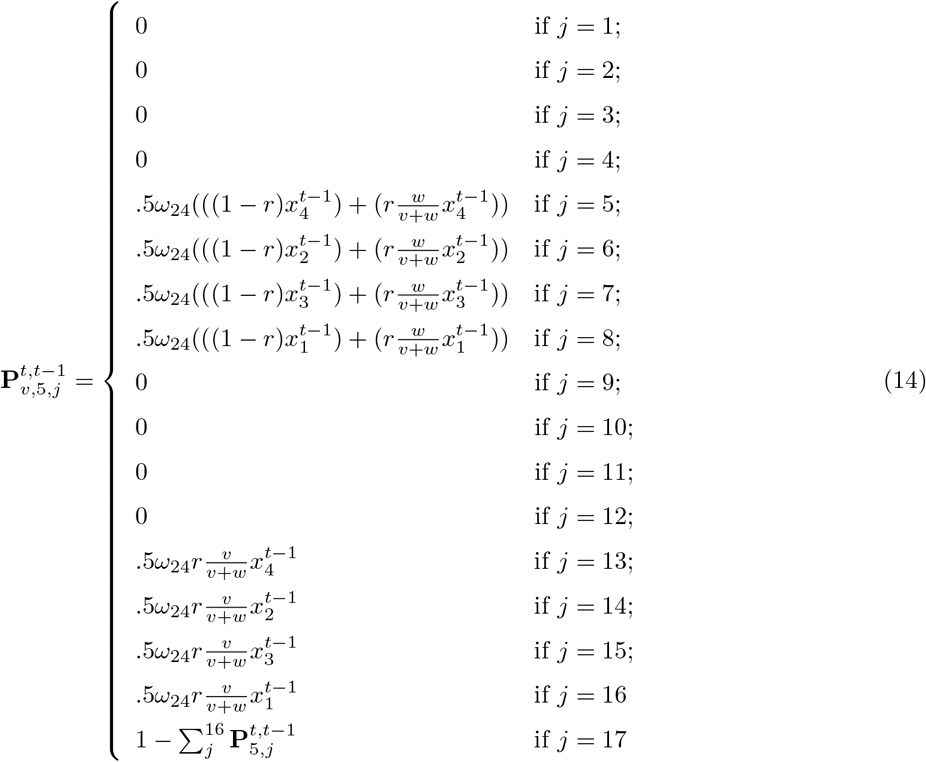

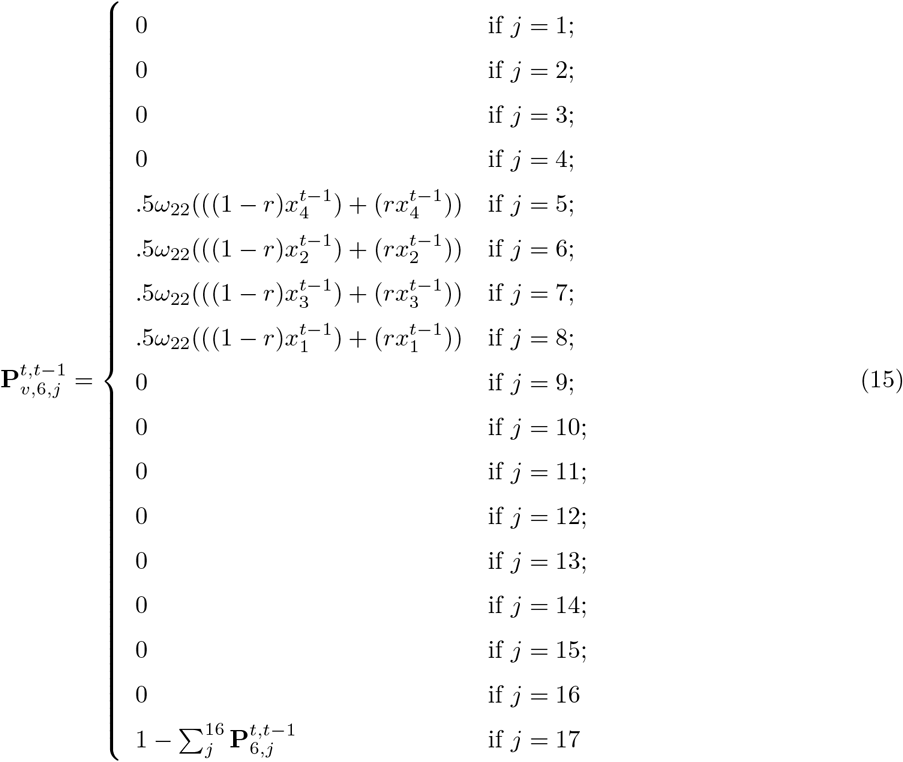

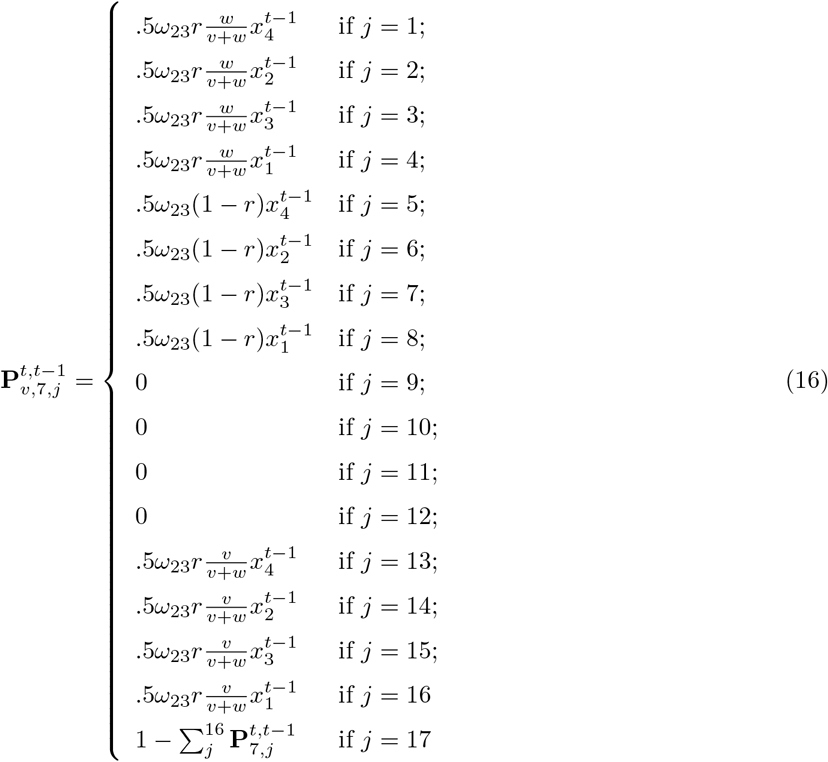

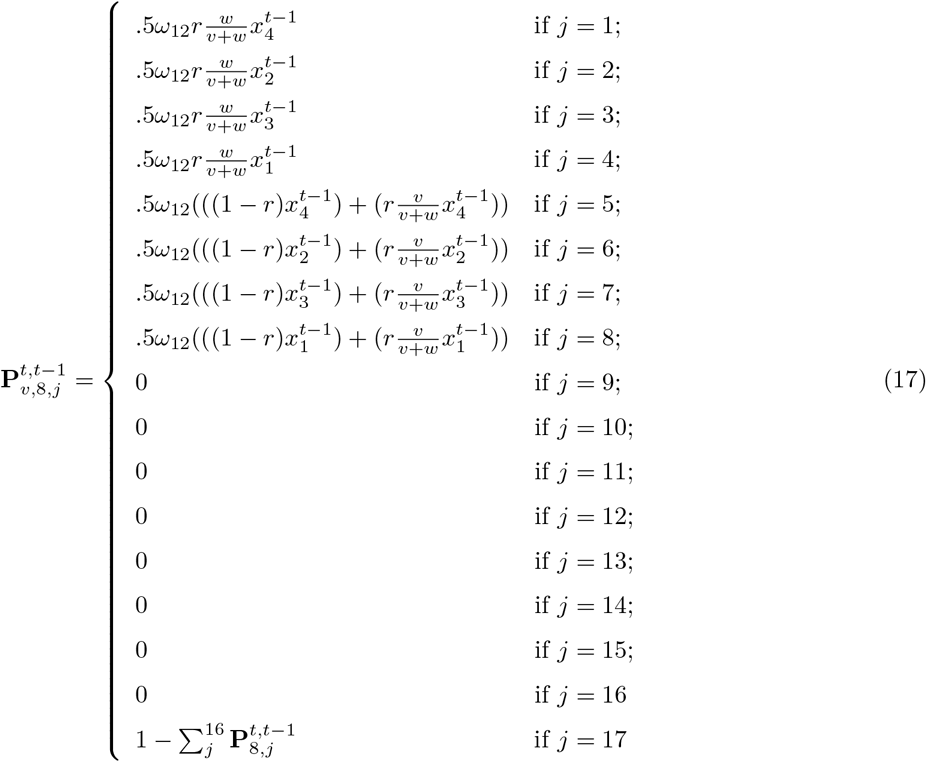

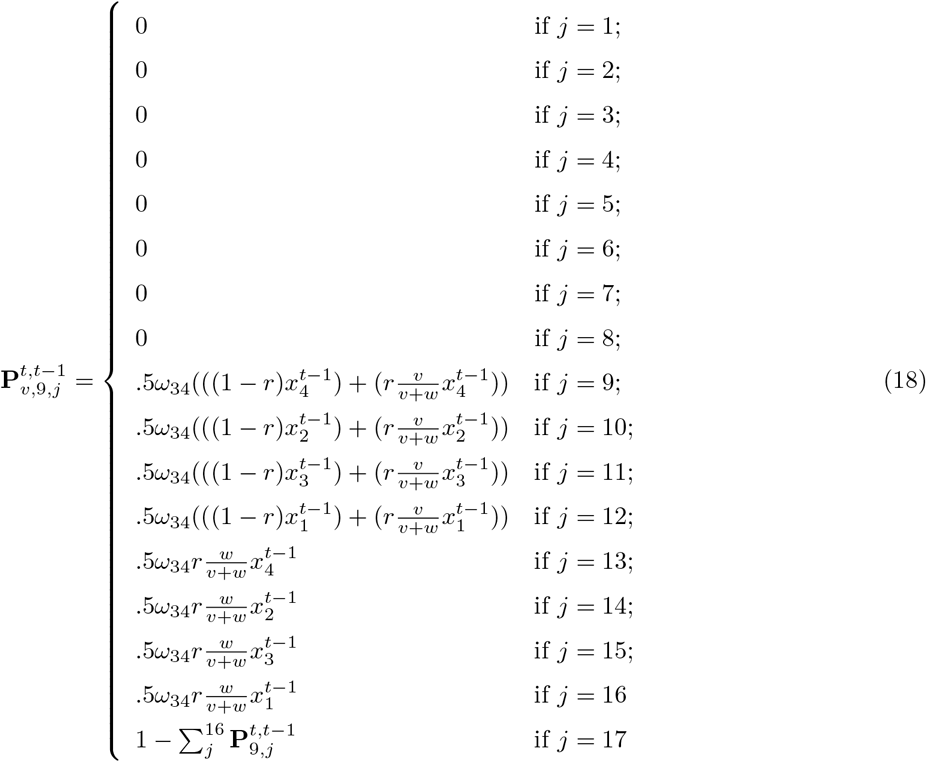

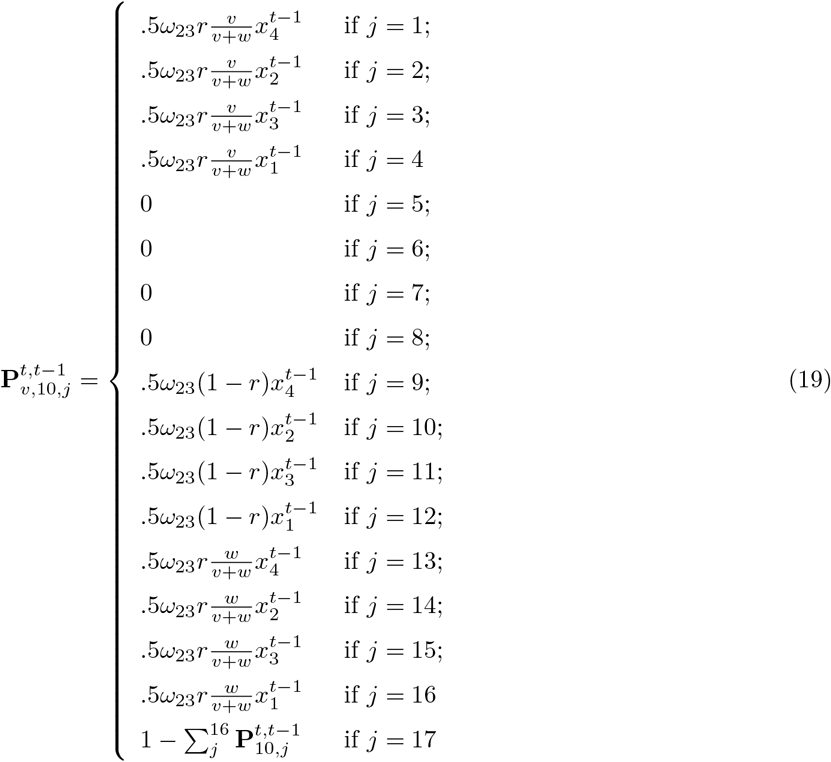

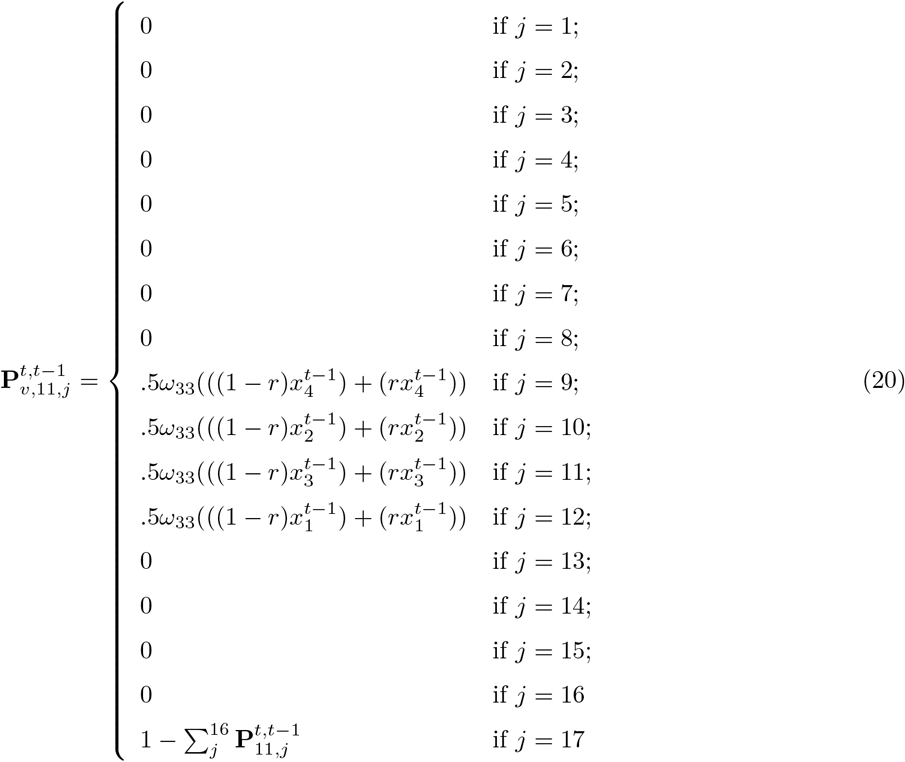

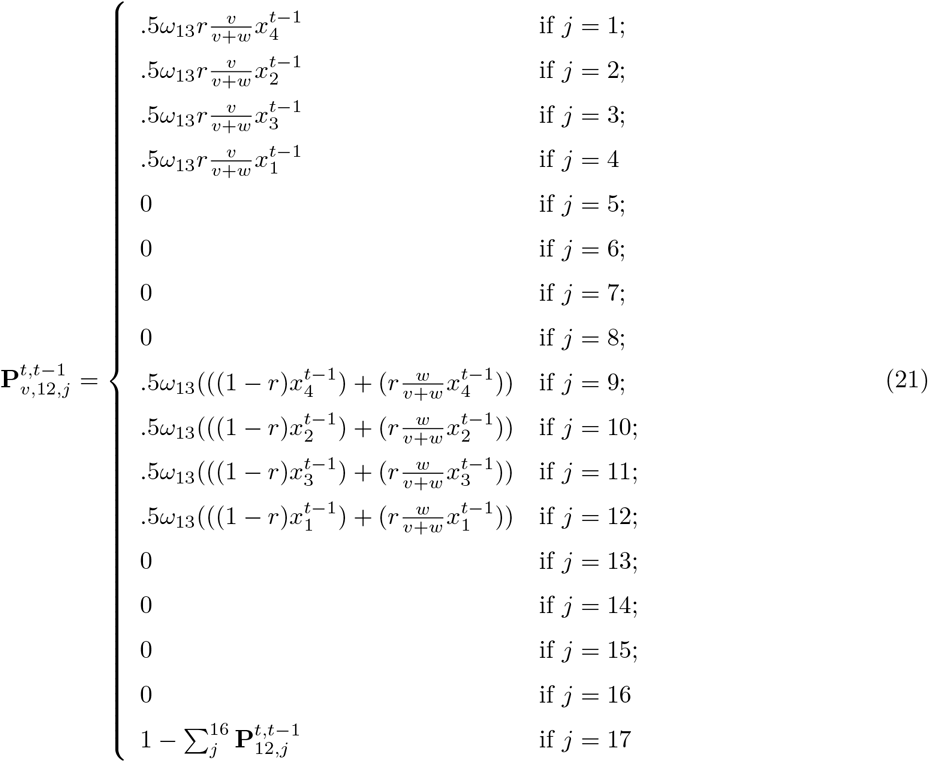

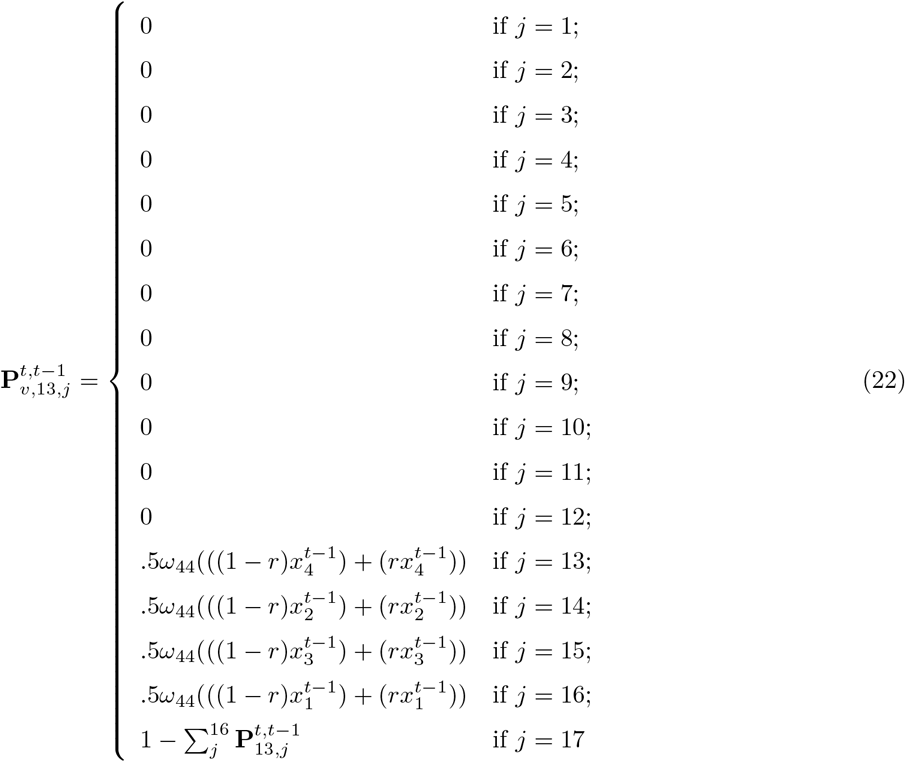

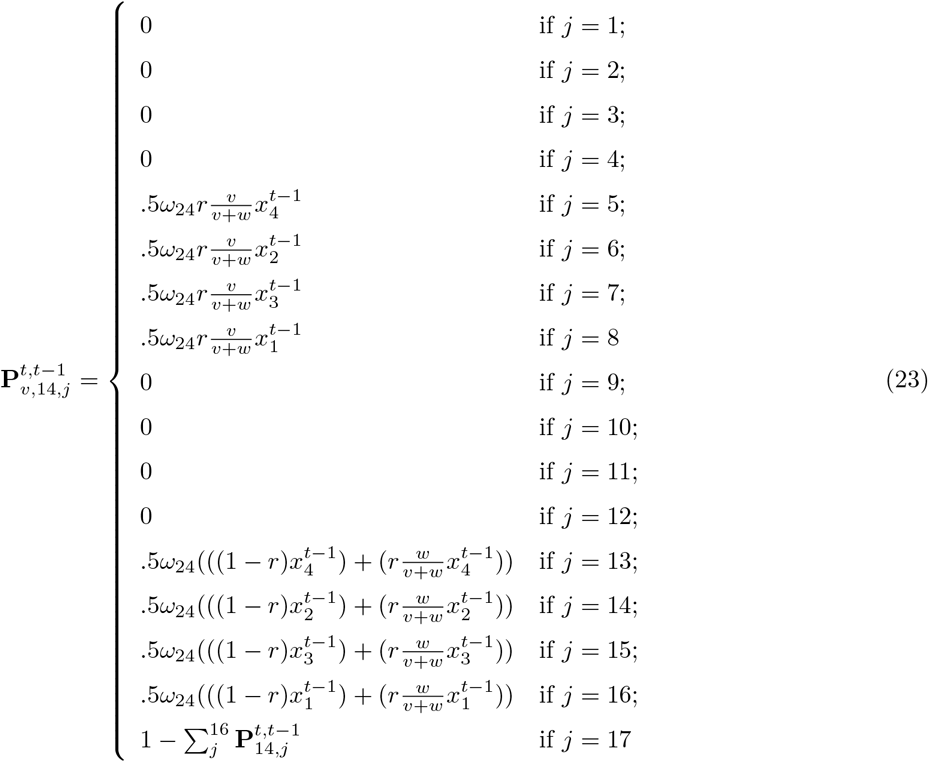

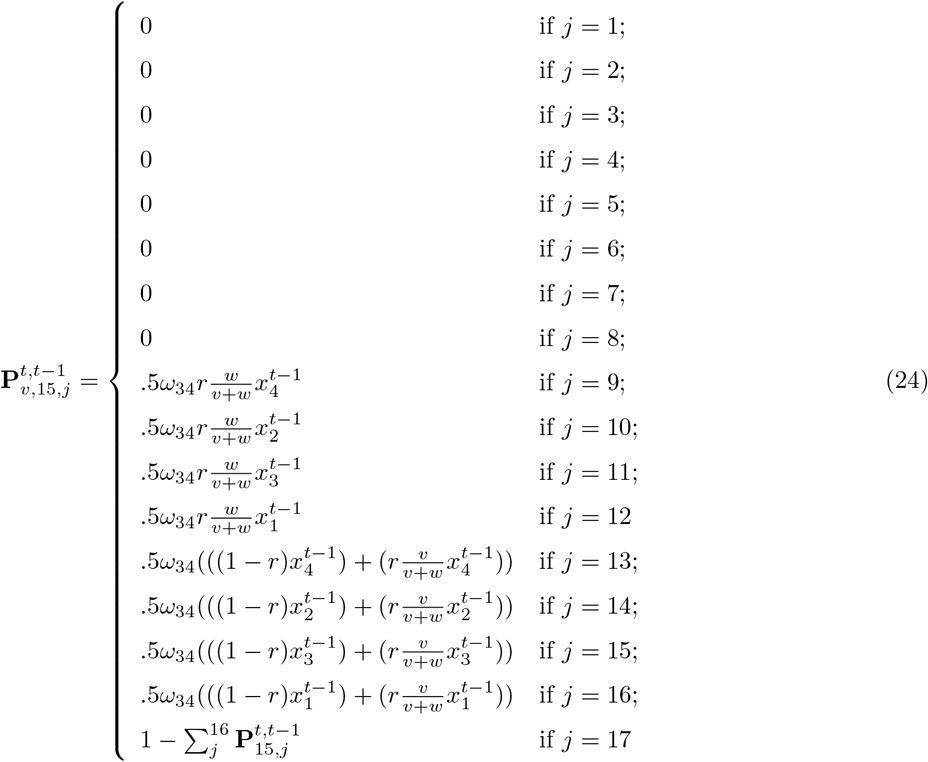

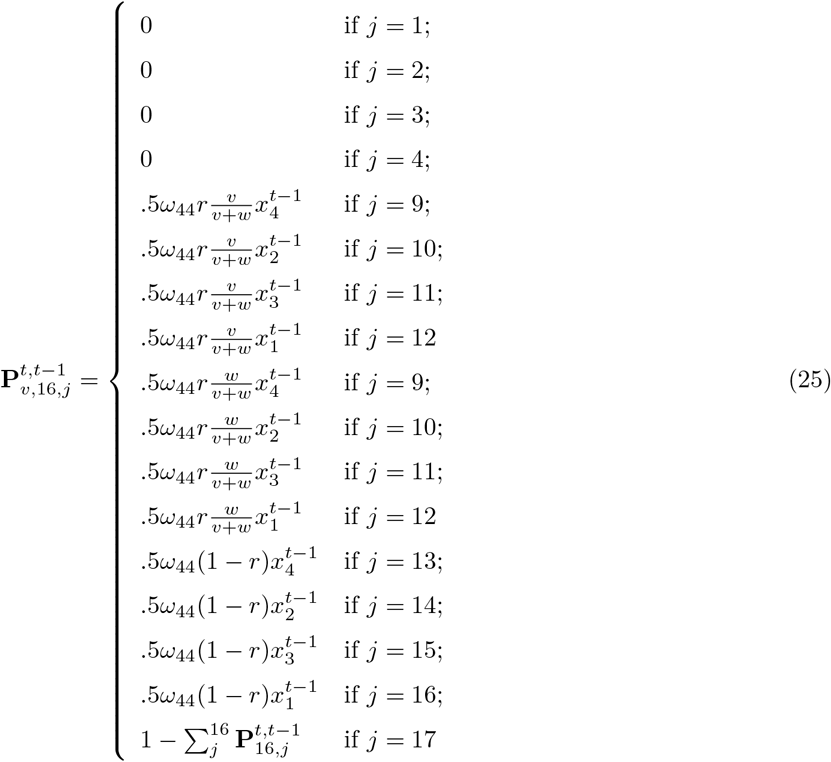

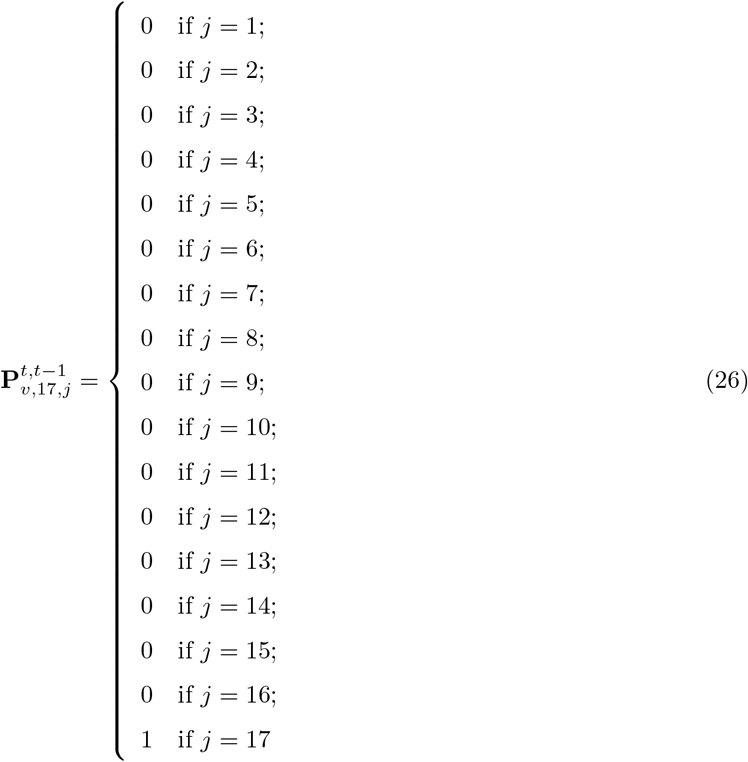

